# Inhibition of *haao-1* enhances oxidative stress response by activating hormetic redox signaling in *C. elegans*

**DOI:** 10.1101/2023.02.16.528568

**Authors:** Raul Castro-Portuguez, Kayla M. Raymond, Emma Thullen, Alana M. Hendrickson, Samuel Freitas, Bradford Hull, Jeremy B. Meyers, Niall Thorns, Emily A. Gardea, Hope Dang, Luis S. Espejo, George L. Sutphin

## Abstract

3-hydroxyanthranilate 3,4-dioxygenase (HAAO) is an intermediate enzyme in the conversion from tryptophan (TRP) to nicotinamide adenine dinucleotide (NAD^+^) via the kynurenine pathway. The kynurenine pathway is the sole *‘de novo’* NAD^+^ biosynthetic pathway from ingested tryptophan. Inhibition of several enzymatic steps in the kynurenine pathway increases lifespan in *Drosophila melanogaster, Saccharomyces cerevisiae*, and *Caenorhabditis elegans*. Knockout or knockdown of *haao-1*, the *C. elegans* gene encoding HAAO, or supplementation of its substrate metabolite 3-hydroxyanthranilic acid (3HAA), has been shown to promote healthy lifespan extension; however, the underlying mechanism remains unknown. In the present study, we report that *haao-1* knockdown induces oxidative stress resistance against several reactive oxygen species (ROS) inducing agents by activating the Nrf2/SKN-1 oxidative stress response pathway. An examination of the redox state of animals with reduced *haao-1* suggests that activation of the Nrf2/SKN-1 pathway is mediated by shifting the balance toward generation of ROS, generating a hormetic effect. Our results identify a novel mechanism for an endogenous metabolite (3HAA) that activates the oxidative stress response. These results provide a conceptual basis by which modulation of the kynurenine pathway can promote healthy aging and enhanced stress resistance.

**Graphical Abstract:** 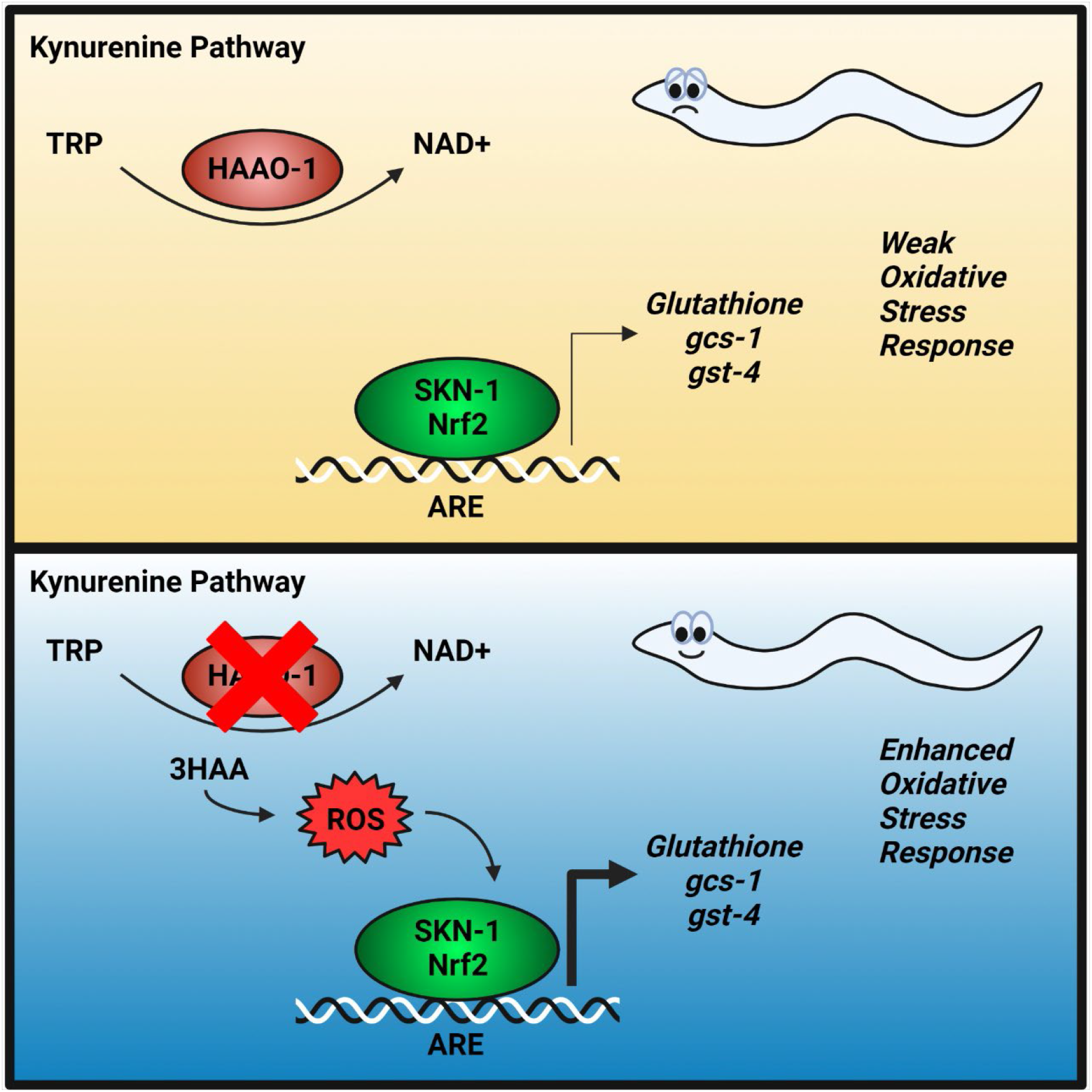

**Highlights:**

- Knockdown of *haao-1* promotes oxidative stress resistance.
- Knockdown of *haao-1* activates the Nrf2/SKN-1 oxidative stress response.
- The shift in redox balance in *haao-1* knockout animals suggests a hormetic mechanism for oxidative stress resistance.

## Introduction

Stochastic accumulation of reactive oxygen species (ROS) and the damage caused by them throughout the entire lifespan has been a prominent theory of causal aging for several decades, however, scientific evidence in the recent years have shown that ROS might be necessary for the normal function of the cell (Beckman and Ames, 1998; Sanz and Stefanatos, 2008; Wickens, 2001). Indeed, exposure to low-level ROS can have a beneficial effect on health span and longevity in several model organisms, a phenomenon termed hormesis (Cypser and Johnson, 2002; Heidler et al., 2010; Przybysz et al., 2009; Yang and Hekimi, 2010). Hormesis is described as a biphasic dose response relationship in which exposure to a small amount of a toxin or harmful agent exerts a beneficial effect in an organism. The mode of action of which a harmful molecule can cause beneficial effects in an organism relies on the low exposure, via minimal dose or transient exposure, to the signal that produces as a compensatory response. This response has been observed as an increased resistance to a subsequent minimal dose of the same stressor or a different stressor (Cypser and Johnson, 2002).

Regulation of several metabolic pathways and endogenous metabolites have shown to upregulate stress responses and mediate beneficial effects against secondary stressors. Itaconate, a mitochondrial metabolite derived from cis-aconitate in the TCA cycle, is required for the activation of the Nrf2 response pathway by LPS in macrophages, playing an important role as regulator of macrophage function and anti-inflammatory (Mills et al., 2018). α-ketoglutarate, another intermediate metabolite in the TCA cycle, is a substrate for prolyl hydroxylase domain-containing enzymes (PHDs), a negative regulator of the hypoxia inducible factor 1α (HIF1α) (Majmundar et al., 2010; Sudarshan et al., 2009).

The kynurenine pathway is a highly conserved metabolic pathway from bacteria to mammals, including *Caenorhabditis elegans* (roundworms), *Drosophila melanogaster* (fruit fly), *Danio rerio* (zebrafish), *Mus musculus* (mouse), and *Homo sapiens* (humans). Its primary function is to convert the ingested tryptophan to nicotinamide adenine dinucleotide (NAD^+^)(Castro-Portuguez and Sutphin, 2020). Cumulative evidence in recent years has implicated dysregulation of this metabolic pathway to a series of stress-related diseases including renal cancer (Hornigold et al., 2020), colorectal cancer (Liu et al., 2021), hepatocellular carcinoma (Shi et al., 2022), neurodegeneration (Campbell et al., 2014; Mor et al., 2021; Schwarcz et al., 2012), and aging (NABEC/UKBEC Consortium et al., 2015; Sutphin et al., 2017). 3-hydroxyanthranilate 3,4-dioxygenase (HAAO) is an intermediary enzyme in the kynurenine pathway that metabolizes 3-hydroxyanthanilic acid (3HAA) into 2-amino-3-carboxymuconate semialdehyde (ACMSA), a precursor to NAD^+^ (**Fig.1A**).

Recent evidence has shown that inhibition of several enzymatic steps in the kynurenine pathway can extend the lifespan in several model organisms. Deletion of BNA1, the yeast ortholog of the HAAO enzyme, can extend chronological lifespan in *Saccharomyces cerevisiae* when grown in 2% glucose (Smith, Jr et al., 2007). A microarray-based genetic screen identified a BNA2, the yeast ortholog of IDO/TDO, mutant to have extended chronological lifespan (Matecic et al., 2010). Oxenkrug showed that genetic (2010) or pharmacological inhibition (2013) of TDO (encoded by the vermillion gene) extended lifespan in the fruit fly *Drosophila melanogaster*. Knockdown of *haao-1*, the gene encoding HAAO in *C. elegans*, or two other gene encoding kynurenine pathway enzymes, *kynu-1* and *tdo-2*, increases in lifespan (Dang et al., 2021; Sutphin et al., 2017; van der Goot et al., 2012). Our initial report on the benefits of *haao-1* knockdown in aging suggested the importance of activating the Nrf2/SKN-1 oxidative stress response, but the specific mechanism of how this happens has not been fully elucidated (Dang et al., 2021).

In this work, we propose that *haao-1* knockdown can increase oxidative stress resistance by increasing the endogenous ROS production and consequently priming the redox balance of the cell to better handle a secondary stressor. First, we show that animals with reduced *haao-1* activity are resistant to multiple forms of oxidative stress. This effect is not dependent on only one of the main transcription factors responsible for handling stress response pathways (*skn-1, daf-16, hif-1* and *hsf-1*). *haao-1* mutants exhibit an upregulation of the Nrf2/SKN-1 antioxidant response that is maintained throughout their lifespan. Finally, we determined that the high antioxidant response of *haao-1* mutants is dependent on the increased of endogenous ROS that primes a hormetic effect in these worms and leads to a higher response to oxidative stressors. Thus, transcriptional factor activation of the main response pathways may cooperate with each other to activate an overall response to stress. Overall, our work demonstrates the role of kynurenine pathway and its intermediate metabolites as signaling molecules to prime the cell against reactive oxygen species and induce a better response to oxidative damage.

## Materials and Methods

### Strains

The following strains were obtained from the Caenorhabditis Genetic Center (CGC) at the College of Biological Sciences at the University of Minnesota: *daf-16(mu86)* I (CF1038), *hif-1(ia4)* V (ZG31), *skn-1(zj15)* IV (QV225) (Tang et al., 2016), *hsf-1(sy441)* I (PS3551), *wgIs341 [skn-1::TY1::EGFP::3xFLAG + unc-119(+)]* (OP341) (Sarov et al., 2006; Zhong et al., 2010), *dvIs19 [(pAF15)gst-4p::GFP::NLS]* (CL2166), *ldIs3 [gcs-1p::GFP + rol-6(su1006)]* (LD1171) (Wang et al., 2010), *mgIs72 [rpt-3p::GFP + dpy-5(+)]* (GR2183), *opIs206 [hif-1p::hif-1(genomic)::GFP::hif-1 3’UTR + unc-119(+)]* (WS4274) (Sendoel et al., 2010), *nIs470 [cysl-2p::GFP + myo-2p::mCherry]* (DMS640), *iaIs7 [nhr-57p::GFP + unc-119(+)]* (ZG120) (Shen et al., 2006), *drSi13 [hsf-1p::hsf-1::GFP::unc-54 3’UTR + Cbr-unc-119(+)]* (OG532) (Morton and Lamitina, 2013), *zcIs4 [hsp-4::GFP]* (SJ4005), *zcIs9 [hsp-60::GFP + lin-15(+)]* (SJ4058), *dvIs70 [hsp-16*.*2p::GFP + rol-6(su1006)]* (CL2070), *muIs84 [(pAD76) sod-3p::GFP + rol-6(su1006)]* (CF1553), *ldIs7 [skn-1b/c::GFP + rol-6(su1006)]* (LD1) (An and Blackwell, 2003), *jrIs1[rpl-17p::HyPer + unc-119(+)]* (JV1) (Back et al., 2012), jrIs2 *[rpl-17p::Grx1-roGFP2 + unc-119(+)]* (JV2) (Back et al., 2012). Strain *haao-1(tm4627)* V (FX04627; backcrossed 6x to N2 to create strain GLS130) were obtained from the C. elegans National Bioresource Project (NBRP) at the School of Medicine at the Tokyo Women’s Medical University. Wild-type (N2) worms were originally obtained from Dr. Matt Kaeberlein (University of Washington, Seattle, WA, USA). Strain OP341 was constructed as part of the Regulatory Element Project, part of modENCODE40. Strain LZR01 *[fmo-2p::mCherry::unc-54]* was gifted by Dr. Scott F. Leiser (University of Michigan, Ann Arbor, MI, USA). Strain haao-1(tm4627)V; wgIs341 (GLS317) was generated by crossing GLS130 to OP341. Strain *haao-1(tm4627)* V; *ldIs3 [gcs-1p::GFP + rol-6(su1006)]* (GLS333) was generated by crossing GLS130 to LD1171. Strain *haao-1(tm4627)* V; *ldIs7[skn-1b/c::GFP + rol-6(su1006)]* (GLS334) was generated by crossing GLS130 to LD1.

### Media and culture

We maintained worms on solid nematode growth media (NGM) seeded with *Escherichia coli* bacteria at 20°C as previously described (Sutphin and Kaeberlein, 2009) except where otherwise noted. RNAi experiments used *E. coli* strain HT115, while all other experiments used strain OP50. All worms were transferred to NGM plates containing 50 μM 5-fluorodeoxyuridine (FUdR) starting at the L4 larval stage to prevent reproduction. Worms were age-synchronized via bleach prep and transferred to plates containing 50 μM 5-fluorodeoxyuridine (FUdR) to prevent reproduction at the L4 larval stage as previously described (Sutphin and Kaeberlein, 2009).

### RNA interference (RNAi)

We conducted RNAi feeding assays according to standard protocols (Sutphin and Kaeberlein, 2009). Briefly, RNAi feeding bacteria were obtained from the Ahringer *C. elegans* RNAi feeding library (Fraser et al., 2000; Kamath et al., 2003). All RNAi plasmids were sequenced to verify the correct target sequence. All experiments were conducted on NGM containing 1 mM Isopropyl β-D-1-thiogalactopyranoside (IPTG) to activate production of RNAi transcripts and 25 μg/mL carbenicillin to select RNAi plasmids and seeded with live *E. coli* (HT115) containing RNAi feeding plasmids.

### Acute juglone toxicity assay

To assess juglone (5-hydroxy-1,4-naphthoquinone, EMD Millipore Cat. No.: 420120-250MG) toxicity, survival assays were conducted as described by Senchuk et al. (2017) Briefly, plates were poured fresh on the day of the assay. To estimate 10-hour survival curves, juglone was prepared as 12 mM stock solution in 100% ethanol while NGM was being autoclaved. Plates were poured with a final concentration of 250 μM juglone, then left to solidify completely in a hood for approximately 15-30 minutes, and spotted with OP50 except where otherwise noted, then left to dry for approximately 30-60 minutes prior to use. Approximately 50 L4 larva were placed on each plate. Experimental animals were scored every hour for 10 hours and counted as dead when they failed to respond to head prodding under a dissection microscope. For 16-hour percent survival assays, the protocol was adjusted to 200 μM juglone and worms were left overnight (approximately 18 hours) and scored for survival the next morning at a single time point.

### Acute paraquat toxicity assay

To assess paraquat (Methyl viologen 98%, Thermo Fisher Scientific Cat. No.: 227320050) toxicity, survival assays were conducted as described by Possik et al.(Possik and Pause, 2015). Briefly, 5-10 synchronized L4 larva were transferred to each well of a 96-well plate containing 50 μL of M9 per well. Then, 50 μL of 100 mM paraquat dissolved in M9 solution were added to each well for a final paraquat concentration of paraquat of 50 mM. Plates were agitated each hour to mix liquid. At least 12 wells per condition were monitored every 4 hours, at which time animals were scored as dead if they were not moving as observed under a dissection microscope.

### Acute arsenic toxicity assay

To assess arsenic (sodium m-arsenite, NaAsO_2_, Sigma-Aldrich Cat. No.: S-7400) toxicity, survival assays were conducted by pouring plates with sodium m-arsenite at a final concentration of 5 mM. Then, plates were spotted with concentrated OP50 bacteria. Approximately 50 L4 larval stage worms were placed on each plate. Experimental animals were scored after 24 hours and counted as dead when not responding to prodding under a dissection microscope.

### Endogenous biosensors assays

Age-synchronized populations of *C. elegans* were maintained as described above. At the designated ages, 15-20 animals were manually transferred to a drop of 25 mM levamisole on slides prepared with 2.2% agarose pads and imaged in triplicate using a Leica M205 FCA Fluorescent Stereo Microscope equipped with a Leica K6 sCMOS monochrome camera using 5x lens (data presented is from at least 30 individual worms per condition). Images were taken using the UV channel (excitation ∼360/40 nm, emission 420 nm) and GFP channel (excitation ∼425/60 nm, emission 480 nm). Briefly, images were saved as LOF files and then exported into RGB tiff files (2048×2048 pixels). Images were first converted to 24-bit RGB tiff files, background was subtracted using automated object segmentation and background values were set to zero. Subsequently, 420-nm images were divided by 480-nm images pixel by pixel to create the ratio images and displayed in false colors using the lookup table “Fire”. Relative H_2_O_2_ levels or GSSG/GSH ratio in these processed images were quantified by using the MATLAB LightSaver script developed in our lab to obtain individual worm data, ratio per pixel histogram compilation and area-under-the-curve (AUC) measurements were extracted per single worm.

### Fluorescence reporter assays

Age-synchronized populations of *C. elegans* were maintained as described above. At the designated ages, 15-20 animals were manually transferred to a drop of 25 mM levamisole on slides prepared with 2.2% agarose pads and imaged using a Leica M205 FCA Fluorescent Stereo Microscope equipped with a Leica K6 sCMOS monochrome camera using 5x lens by triplicate (data presented is from at least 30 individual worms per condition). Identical imaging settings were maintained across timepoints for each strain. Images were not processed and then exported as TIFF files using the acquisition software LAS X. Data analysis was performed using the MATLAB LightSaver script developed in our lab to obtain individual worm data, intensity per pixel histogram compilation and area-under-the-curve (AUC) measurements were extracted per single worm. Each experiment was repeated in biological duplicate.

### ROS measurements

For 2′,7′-dichlorodihydrofluorescein diacetate (H_2_DCFDA, Invitrogen(tm) Cat. No.: D399). After treatment, approximately 50 worms per condition were washed from the agar plate using M9 buffer and washed three times in M9 buffer. Worms were kept in a conical tube with 3mL of M9 and H_2_DCFDA was added to each tube at a final concentration of 25 μM. Tubes were kept wrapped in aluminum foil at room temperature for 4 hours while being in a nutator. Then, worms were transferred to a NGM plate spotted with OP50 to allow them crawl for a couple of minutes. 15-20 animals were manually transferred to a drop of 25 mM levamisole on slides prepared with 2.2% agarose pads and imaged using a Leica M205 FCA Fluorescent Stereo Microscope equipped with a Leica K6 sCMOS monochrome camera using 5x lens by triplicate.

### Data Analysis

All graphs were made using GraphPad Prism software (version 8. 4.3.686). P-values for statistical comparison of fluorescence fold change between test groups were calculated by parametric unpaired t-test (two-sided) with Welch’s correction and using the Holm-Sidak multiple test correction method, with alpha = 0.05. Significance in P values are notated with asterisks as it follows: * if p < 0.05, ** if p < 0.01, *** if p < 0.001, and **** if p < 0.0001. Most plots show individual data points while the bar graphs represent mean value, and the error bars represent the standard error of the mean (SEM). For survival curves assays, the statistical groupwise and pairwise comparisons among survivorship curves were performed by. P values were obtained using the log-rank analysis (select pairwise comparisons and group comparisons or interaction studies) as noted. Summary survival data, sample size (n), and statistics are included in Supplementary Data 2.

## Results

### Inhibition of *haao-1* improves resistance to oxidative stress in *C. elegans*

Our lab first showed that *haao-1* knockdown and knockout, as well as treatment with 1mM 3HAA, increases survival of *C. elegans* challenged with 20 mM paraquat. As an initial step, we sought to examine the oxidative stress resistance of worms with reduced *haao-1* expression to a variety of oxidative stressors. We examined the sensitivity of homozygous *haao-1* deletion worms (*haao-1(tm6427)*) (**Fig. 1B**) or wild type worms treated with *haao-1(RNAi)* to a variety of oxidative stress inducers: juglone 250 μM (**Fig. 1C**,**D**), paraquat 50 mM (**Fig. 1E**,**F**) and arsenic 5 mM (**Fig. 1G**). In each case, the *haao-1* mutants or RNAi treated worms (**Fig. 1B**) exhibited an increased survival. Our observation that *haao-1* inhibition promotes resistance to agents that induce oxidative stress through distinct mechanism supports a model in which *haao-1* inhibition removes ROS and does not specifically inhibit the distinct mechanism or subcellular compartment of ROS generation (e.g., superoxide generated by inhibition of mitochondria electron transport chain complex). Because the degree of resistance induced by *haao-1* inhibition varies across the different stress inducing agents, *haao-1* may be better suited to handle specific subspecies of ROS, specific forms of damage, or also beneficially interact with other molecular processes such as proteasomal degradation, proteotoxic stress, and osmotic stress.

**Figure 1.**
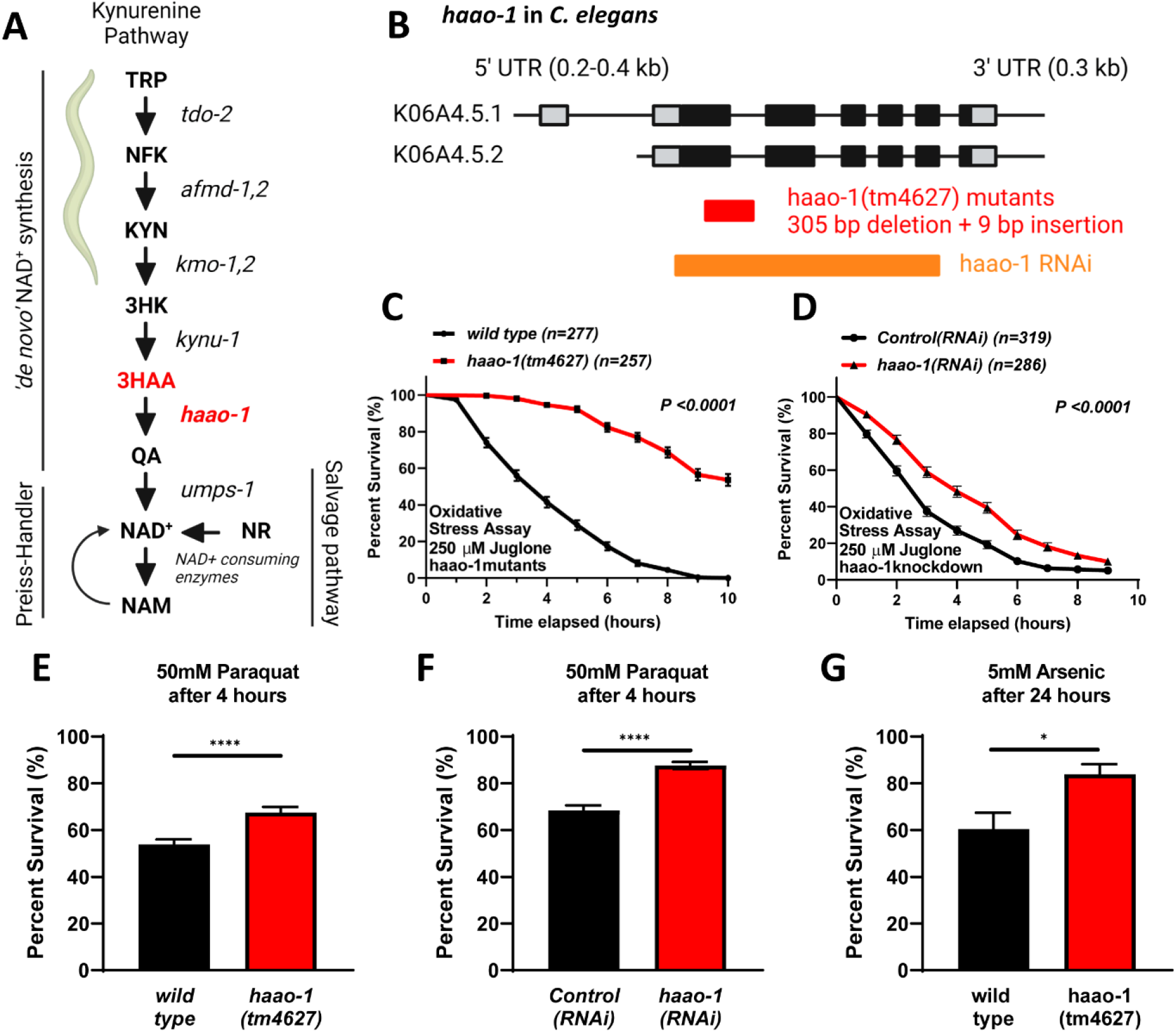
haao-1inhibition enhances oxidative stress resistance in C. elegans. (A) Scheme of the kynurenine pathway, its enzymes and metabolites. (B) Genomic organization of the K06A4.5 (*haao-1*) locus (gray is untranslated UTR; black are translated exons; adapted from wormbase.org). The *haao-1* gene encodes two isoforms (K06A4.5.1 and K06A4.5.2). The tm4627 allele (red) is a 305 bp deletion with 9bp insertion. RNAi clone (orange) is from Ahringer RNAi library. (C) Survival of wild type (N2) vs *haao-1(tm4627)* mutants or (D) wild type (N2) in empty vector (*control(RNAi)*) vs *haao-1(RNAi)* at L4 stage in 250 μM juglone. P values were determined by log-rank. (E) Survival of wild type (N2) (n=720) vs *haao-1(tm4627)* mutants (n=711) or (F) wild type (N2) in empty vector (*control(RNAi)*) (n=581) vs *haao-1(RNAi)* (n=625) at L4 stage in 50 mM paraquat for 4 hours. (G) Survival of wild type (N2) (n=238) vs *haao-1(tm4627)* mutants (n=243) at L4 stage in 5 mM arsenic for 24 hours. P-values were determined by parametric unpaired t-test with Welch’s correction and using the Holm-Sidak multiple test correction method, with alpha = 0.05. Enzymes: tryptophan 2,3-dioxygenase (tdo-2); arylformamidase (afmd-1,2); kynurenine 3-monooxygenase (kmo-1,2); kynureninase (kynu-1); 3-hydroxyanthranilate 3,4-dioxygenase (haao-1); and uridine monophosphate synthetase (umps-1). Metabolites: tryptophan (TRP); N-formylkynurenine(NFK); kynurenine (KYN); 3-hydroxykynurenine (3HK); 3-hydroxyanthranilic acid (3HAA); quinolinic acid (QA); nicotinamide adenine dinucleotide (NAD+), nicotinamide (NAM); and nicotinamide riboside (NR).

### Central oxidative stress response transcription factors are dispensable for *haao-1*-induced oxidative stress resistance

Several transcription factors upregulate response pathways to handle free radicals in the cell. Previous work has shown that ROS are capable of increasing transcriptional activity of several transcription factors, including *skn-1* (ortholog of the mammalian Nrf2) (Ewald et al., 2017; van der Hoeven et al., 2011), *daf-16* (ortholog of the mammalian FOXO family) (Senchuk et al., 2018; Zou et al., 2013), *hif-1* (ortholog of mammalian HIF1A) (Hwang et al., 2014; Qutub and Popel, 2008), and *hsf-1* (ortholog of the HSF1 transcription factor) (Hartwig et al., 2009). To assess if any of these transcription factors were necessary for the upregulated oxidative stress response upon *haao-1* inhibition, we tested wild type and TF mutant worms under juglone acute exposure for approximately 16 hours. In all the conditions, except in *hsf-1(sy441)* mutants, inhibition of haao-1 using RNAi fed bacteria was able to increase the survival of all the mutant strains in comparison to the control (**Fig. 2A**). We also determined that *hsf-1* animals were already resistant to juglone and were alive at 16 hours, so we were unable to detect an additional increase in resistance for *haao-1(RNAi)* if it was present.

**Figure 2.**
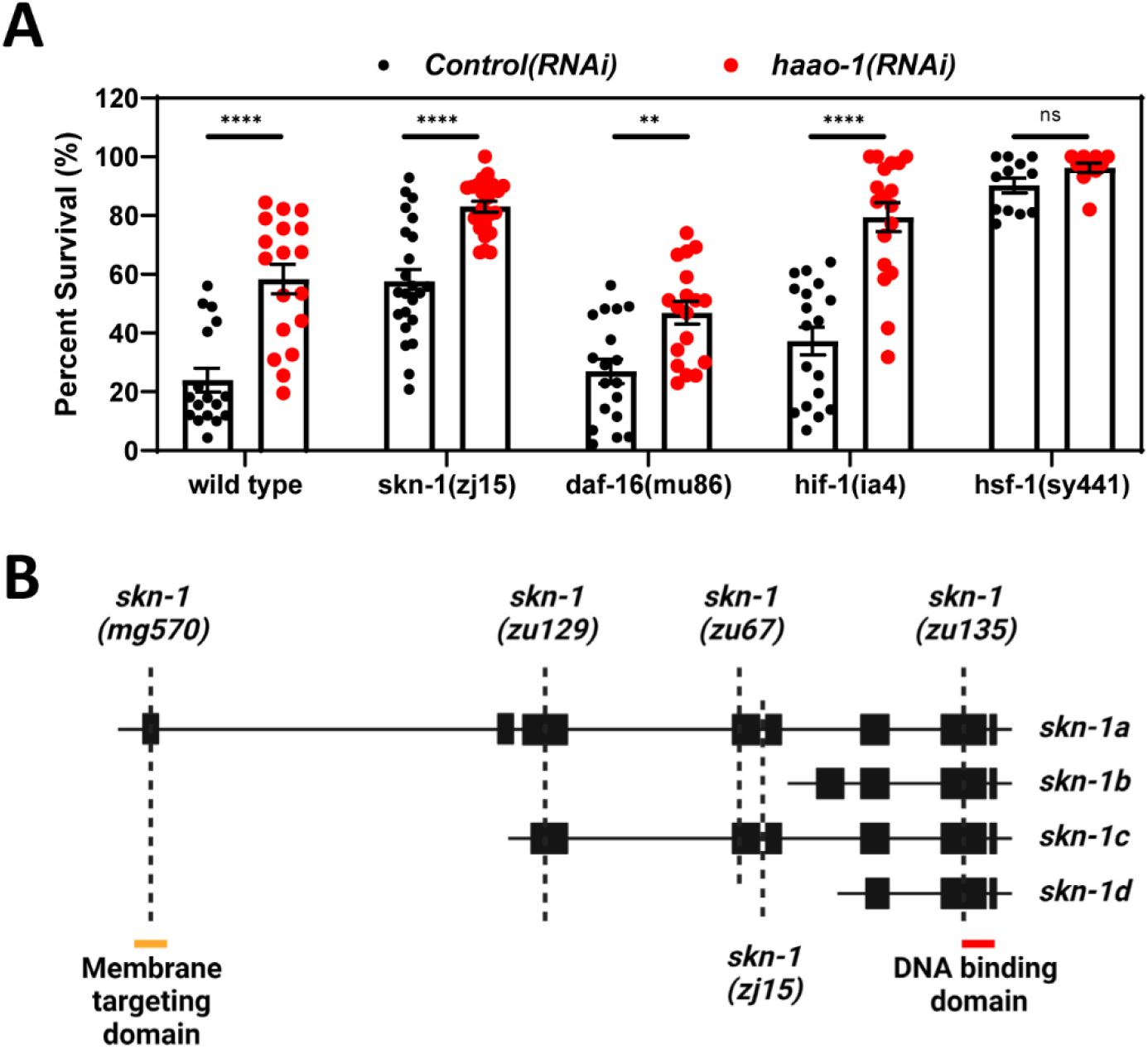
Four main transcription factors are not required for the enhanced oxidative stress response upon haao-1 inhibition. (A) Wild type (N2) and transcription factor mutant worms [*skn-1(zj15), daf-16(mu86), hif-1(ia4)* and *hsf-1(sy441)*] grown from egg in plates containing empty vector (*control(RNAi)*) or *haao-1(RNAi)*. At L4 larval stage they were subjected to 200 μM juglone exposure for 16 hours. Experimental animals were scored and counted as dead when not responding to prodding under a dissection microscope. N values and statistics can be found in Supplementary Data 2. (B) Genomic organization of the T19E7.2 (skn-1) locus (black are translated exons, red is DNA binding domain, yellow is membrane targeting domain; adapted from wormbase.org). P-values were determined by parametric unpaired t-test with Welch’s correction and using the Holm-Sidak multiple test correction method, with alpha = 0.05.

### *haao-1* inhibition induces multiple oxidative stress pathways and this induction is maintained throughout the lifespan

To further explore which pathways are responsible for this stress response, we determined the effect of *haao-1(RNAi)* on transgenic fluorescent reporters in at least 4 major molecular pathways. We observed that *haao-1* knockdown caused a major increase in each stress response pathway, and that this induction was maintained throughout the entire lifespan (**Fig. 3**). *haao-1(RNAi)* induced accumulation of protein levels of the master regulator of the antioxidant response SKN-1.Its transcriptional activity is confirmed by showing increased expression of several well-characterized transcriptional targets: gamma-glutamylcysteine synthetase (*gcs-1*), glutathione S-transferase 4 (*gst-4*), both part of the glutathione (GSH) synthesis and metabolism pathway, as well as 26S proteasome regulatory subunit, *rpt-3* (**Fig. 3A**). Interestingly, *haao-1(RNAi)* treated worms also induce accumulation and protein expression of the HIF-1 transcription factor as well as 3 well-characterized downstream genes, cysteine synthase like-2 (*cysl-2*), nuclear hormone receptor family member 57 (*nhr-57*), and flavin-containing monooxygenase 2 (*fmo-2*) (**Fig. 3B**). The heat shock transcription factor 1 (HSF-1)-regulated response pathway also is active upon *haao-1* inhibition using RNAi and 3 main heat shock proteins (*hsp-4, hsp-60* and *hsp-16*.*2*) are significantly increased in compares with its control counterpart (**Fig. 3C**). We did not observed accumulation of DAF-16 in puncta in the nucleus as has been observed in the case of a heat shock response, however, we observed an increase in the transcriptional activity of one of its downstream targets, superoxide dismutase 3 (*sod-3*) (**Fig. 3D**). These four main response pathways are active upon *haao-1* inhibition using RNAi and they are maintained throughout the lifespan of the nematodes (**Fig. 3F**).

**Figure 3.**
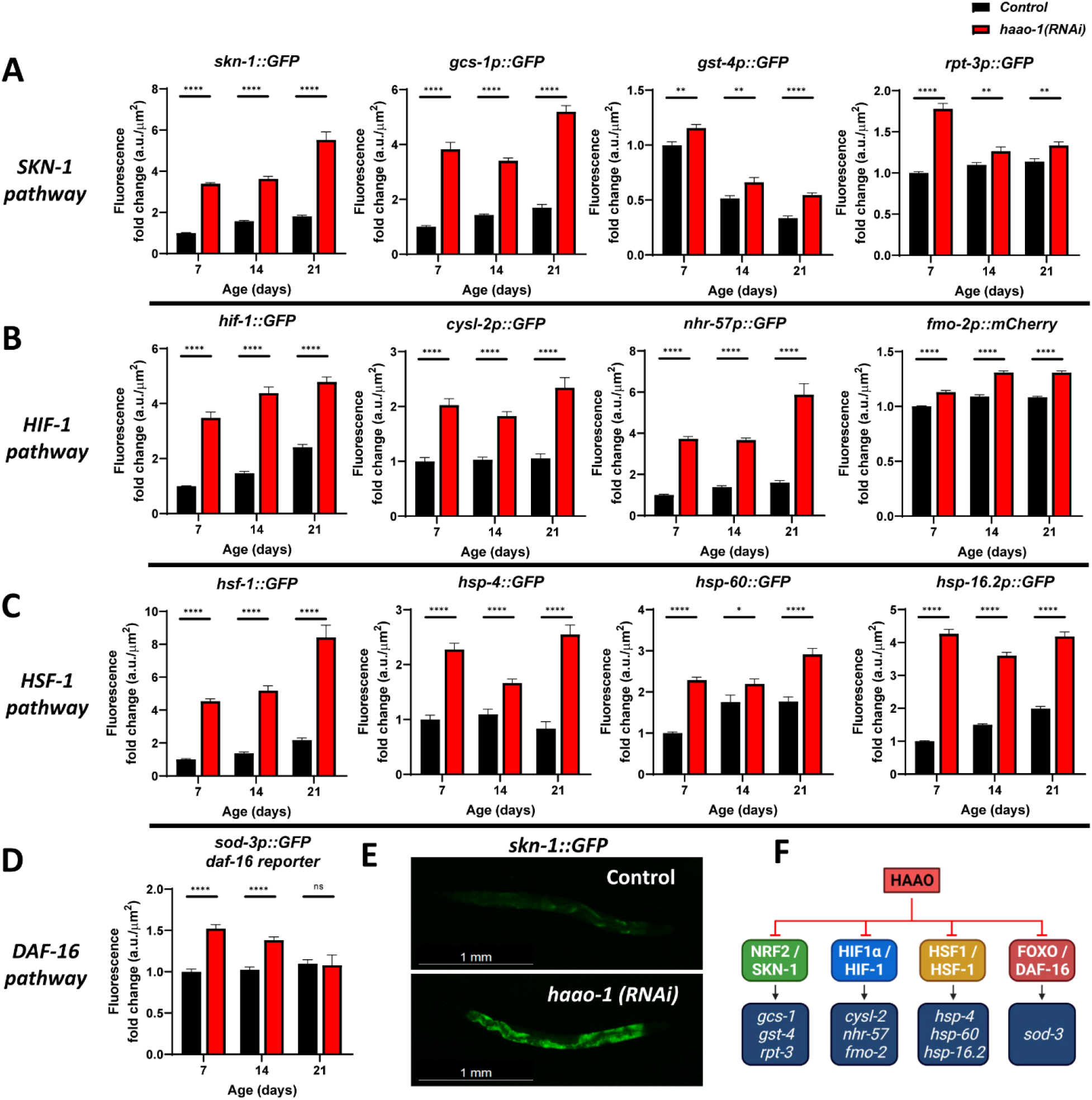
haao-1inhibition induces activation of four main transcription factors responsible for enhanced oxidative stress resistance. (A) GFP expression of SKN-1 transcription factor (ortholog of Nrf2) and its activity by the downstream genes *gcs-1, gst-4*, and *rpt-3*. (B) GFP expression of HIF-1 transcription factor (ortholog of HIF1α) and its activity by the downstream genes *cysl-2, nhr-57*, and *fmo-2*. (C) GFP expression of hsf-1transcription factor (ortholog of HSF1) and its activity by the downstream genes *hsp-4, hsp-60*, and *hsp-16*.*2*. (D) GFP expression of DAF-16 transcription factor (ortholog of FOXO) by its reporter gene *sod-3*. (E) Example of images taken for the *skn-1::GFP* strain in *control(RNAi)* versus *haao-1(RNAi)* at day 7 from egg. Number of worms analyzed per bar is approximately n=30 worms. (F) Representative scheme of the effects caused by haao-1and its inhibition in different main transcription factors. P-values were determined by parametric unpaired t-test with Welch’s correction and using the Holm-Sidak multiple test correction method, with alpha = 0.05.

### Endogenous ROS production is increased upon haao-1 inhibition

We then sought to understand what common causes can upregulate multiple response pathways in *C. elegans*. We asked whether ROS levels were elevated in worms upon *haao-1* inhibition. We first use the *jrIs1[rpl-17p::Hyper]* reporter strain that encodes an endogenous biosensor of ROS inside the worm. We observed a marked increase of ROS levels at day 7 from egg.(**Fig. 4B**) We confirmed this upregulation of ROS using the ROS-sensitive H2DCFDA dye (**Fig. 4D**). Interestingly, the increase in ROS declined with increasing age, with levels actually lower for *haao-1(RNAi)* animals relative to *control(RNAi)* by day 21 from egg. To further observe the impact of ROS we used transgenic worms expressing a biosensor reporting the oxidation state of the endogenous antioxidant glutathione *jrIs2 [rpl-17p::Grx1-roGFP2]*. The oxidized-to-reduced glutathione ratio was higher at day 7 for animals subjected to *haao-1(RNAi)* relative to control and, following the trend for endogenous ROS, also declined with age until day 21. (**Fig. 4C**)

**Figure 4.**
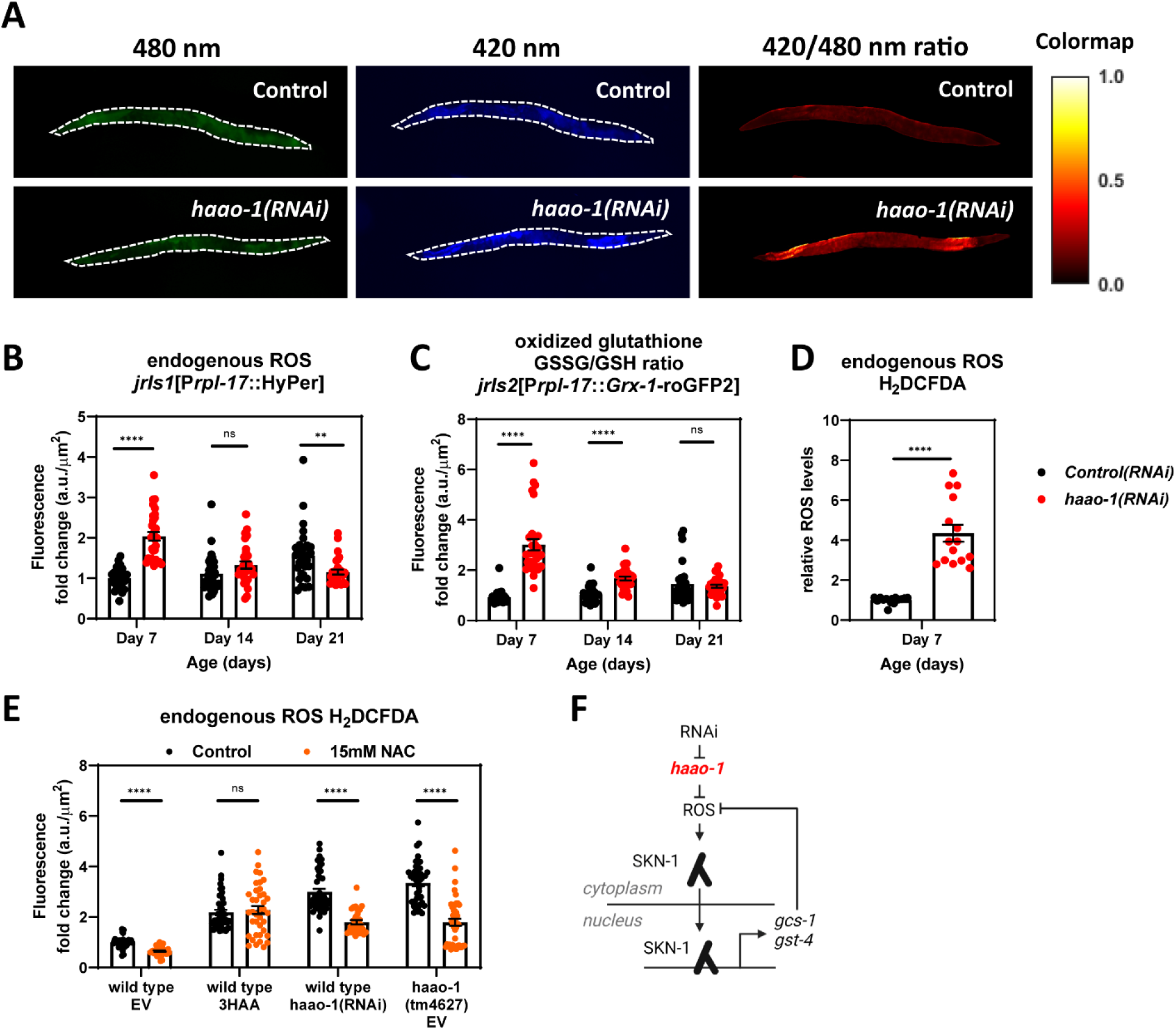
haao-1inhibition induces endogenous production of ROS. (A) Representative images of *jrIs1[rpl-17p::Hyper]* strain on *control(RNAi)* vs *haao-1(RNAi)*. Images taken at 420nm and 480nm emission in the green and blue channels. Pixel-by-pixel ratio was determined and faked-color in the right panels. (B) Endogenous ROS levels were determined using *jrIs1[rpl-17p::Hyper]* strain and (C) Oxidized glutathione ratio was determined using the *jrIs2[rpl-17p::Grx-1-roGFP2]* strain, quantification was made using the LightSaver script. (D) Endogenous ROS were also determined using the fluorescent-dye H2DCFDA in day 7 adult worms exposed to *control(RNAi)* vs *haao-1(RNAi)*. Number of worms analyzed per bar is approximately n=30 worms. (E) Relative endogenous ROS determined by H2DCFDA in day 7 adult worms exposed to control plates or 15mM N-acetylcysteine (NAC) from egg. Number of worms analyzed per bar is approximately n=45 worms. (F) Schematic diagram of *haao-1*-mediated induction of SKN-1 by increased production of ROS. P-values were determined by parametric unpaired t-test with Welch’s correction and using the Holm-Sidak multiple test correction method, with alpha = 0.05.

To further dissect the effects of haao-1 inhibition and increase in ROS levels, we treated worms with N-acetylcysteine (NAC) a glutathione precursor that can reduce endogenous ROS levels. In all the cases, we observed how worms treated with 1mM 3HAA, treated with *haao-1(RNAi)* or *haao-1(tm4627)* have an increased levels of ROS based on the fluorescence of the oxidized DCF dye. In wild type worms, NAC was able to reduce the ROS in a small portion; however, the decrease of ROS was more notorious in the *haao-1(RNAi)* treated worms and in the *haao-1(tm4627)* mutant worms. We did not see a clear effect in the worms treated with 3HAA and this could be due to the fact that 3HAA and its redox properties could interfere with the NAC or vice versa by interacting with each other in the agar plates.(**Fig. 4E**)

### haao-1 inhibition induces activation of the transcription factor skn-1 through ROS-dependent and ROS-independent mechanisms

Canonical activation of skn-1 can be accomplished through either ROS or kinase activation, for instance *pmk-1* (an ortholog of the MAP kinase p38). (Ewald et al., 2017). We next sought to identify which mechanisms were mediating SKN-1 activation in response to *haao-1* knockdown. 15 mM NAC was sufficient to decrease ROS levels in *haao-1* knockout animals back to wild type levels (**Fig 5A**); however, while 15 mM NAC did substantially reduce SKN-1 activity, neither *skn-1* expression nor its transcriptional activity returned to basal levels following 15 mM NAC (**Fig. 5B**,**C**). This suggested that haao-1 inhibition induces skn-1 transcriptional activation via both ROS-dependent and ROS-independent mechanisms.

**Figure 5.**
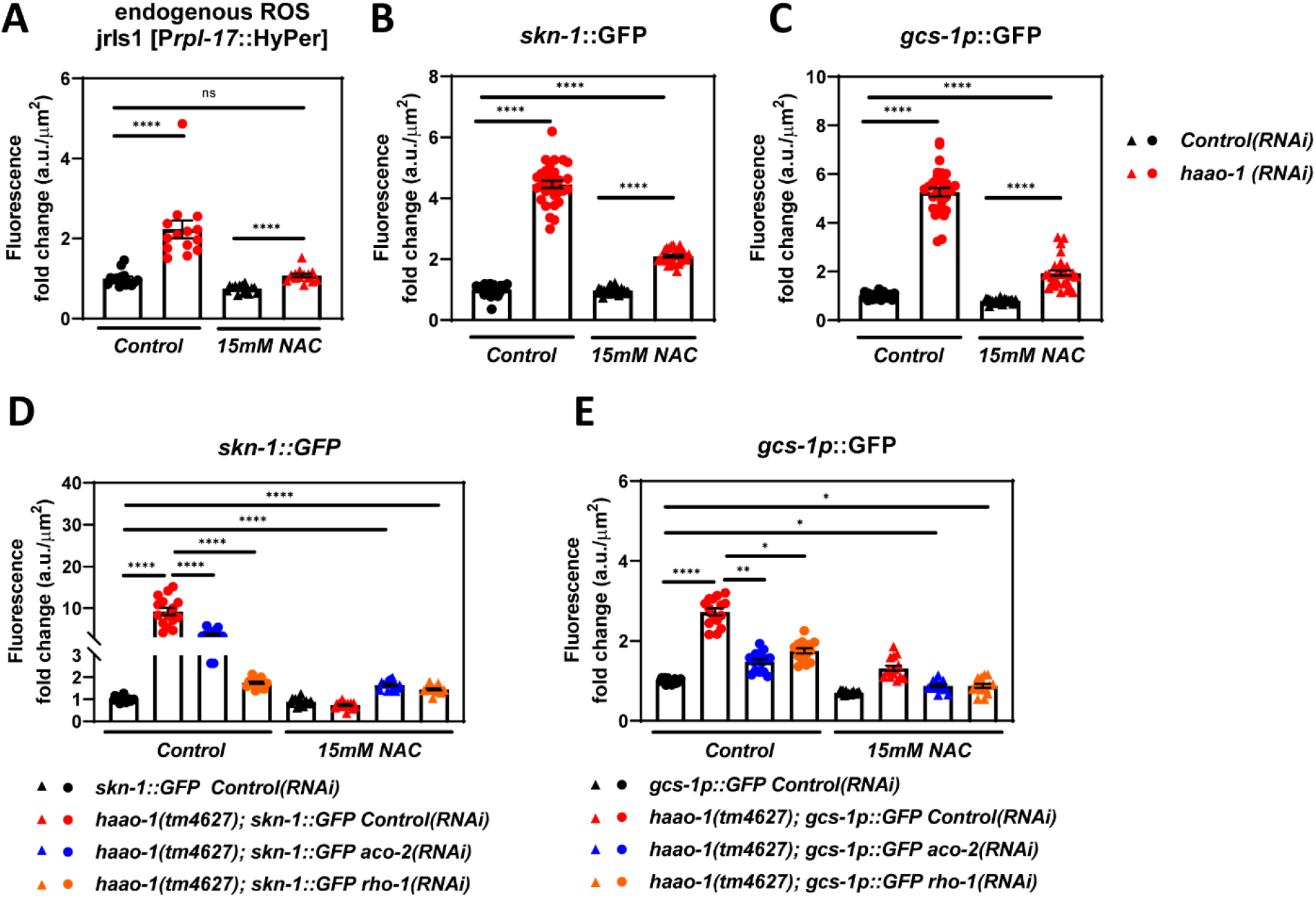
Increased endogenous ROS production in haao-1 knockout background is partially responsible for the activation of skn-1. (A) Endogenous ROS levels were determined using *jrIs1[rpl-17p::Hyper]*strain (B) SKN-1 protein expression levels were determined using a *skn-1::GFP* strain (C) GFP expression driven by *gcs-1* promoter, a downstream gene of *skn-1*(D-E) combination of 15 mM NAC with *aco-2(RNAi)* or *rho-1(RNAi)* can bring *skn-1* expression and *gcs-1* to back basal levels in *haao-1(tm4627)*background. P-values were determined by parametric unpaired t-test with Welch’s correction and using the Holm-Sidak multiple test correction method, with alpha = 0.05.

To identify the mediators of the ROS-independent activation of SKN-1 by *haao-1* knockdown, we used RNAi to screen selected genes for the capacity to reduce the activation of SKN-1 by *haao-1(tm4627)*. The genes selected for the screening (**Supplementary Fig. 2**) were based on potential mediator routes of SKN-1 activation or pathways that are activated in response to several stressors. As we anticipated, of the 85 tested genes, we observed that most RNAi did not reduce the activation of SKN-1 by *haao-1(tm4627)* (**Supplementary Fig. 2**). Of the 85 genes, knockdown of 11 significantly reduced SKN-1::GFP in the *haao-1(tm4627)* background. At this stage, it remained possible that the 11 RNAi identified in the primary screen either (1) reduced SKN-1 activation either by reducing ROS, and thus interacting with the ROS-dependent component of *haao-1* mediated SKN-1 activation, or (2) were mediating the ROS-independent activation of SKN-1 by *haao-1*.

To distinguish between these possibilities, we performed a secondary screen to investigate which of the 11 RNAi could retain the capacity to reduce SKN-1 activation even in the presence of 15 mM NAC. Unlike our earlier experiments with NAC, which used *haao-1(RNAi)*, this secondary screen required the use of *haao-1* to allow the screen to be conducted using RNAi. However, when we crossed the haao-1(tm4627) knockout strain to the *skn-1::GFP* transgenic strain, the elevated expression of skn-1::GFP was completely repressed by 15 mM NAC. (**Supplementary Fig. 3A**). To further explore this potential confounder, we crossed the *haao-1(tm4627)* knockout strain to the SKN-1 transcriptional reporter strain transgenically expressing *gcs-1p::GFP*. (**Supplementary Fig. 3B**) As we observed with *haao-1(RNAi)*, this strain retained residual SKN-1 activity induced by *haao-1(tm4627)* even in the presence of 15 mM NAC (**Supplementary Fig. 3B**). In that context, is clear that NAC is able to diminish skn-1 transcriptional activity in the haao-1(tm4627) background, however, the skn-1 activation is not completely eliminated. We proceeded to conduct our secondary screen using both *haao-1(tm4627)*; *skn-1::GFP* and *haao-1(tm4627)*; *gcs-1p::GFP* reporter strains. Of the 11 genes examined in this secondary screen, RNAi knockdown of two consistently decreased skn-1 transcriptional activity in the context of *haao-1* knockout combined with 15 mM NAC: *aco-2*, a mitochondrial aconitase and ortholog of the human ACO2, and *rho-1*, a RHO family small GTPase and ortholog of human RHOA (**Fig. 5D**,**E**). Combining 15 mM NAC with either *aco-2(RNAi)* or *rho-2(RNAi)* completely ablated the transcriptional activation of SKN-1 by *haao-1(tm4627)*.

### Increased ROS upon haao-1 inhibition is necessary for its resistance to oxidative stress

To test the possibility that ROS production is necessary for *skn-1* activation and oxidative stress resistance in the *haao-1(tm4627)* background, we test survival of wild type and *haao-1* mutant worms in response to juglone in the presence of NAC. NAC was sufficient to increase the glutathione production in wild type worms, giving them an increase in survival that is likely dependent on the ability of the system to upregulate the GSH biosynthetic pathway (**Fig. 6A**). *haao-1(tm4627)* animals were more resistant to the juglone as previously shown using haao-1(RNAi) (**Fig 2A**). Interestingly, when we treated *haao-1(tm4627)* worms with 15mM NAC the survival benefit of the *haao-1(tm2627)* worms challenged with juglone was completely eliminated. (**Fig. 6A**) This supports the model that the endogenous increase in ROS in *haao-1(tm4627)* is necessary for the increased resistance to oxidative stress.

**Figure 6.**
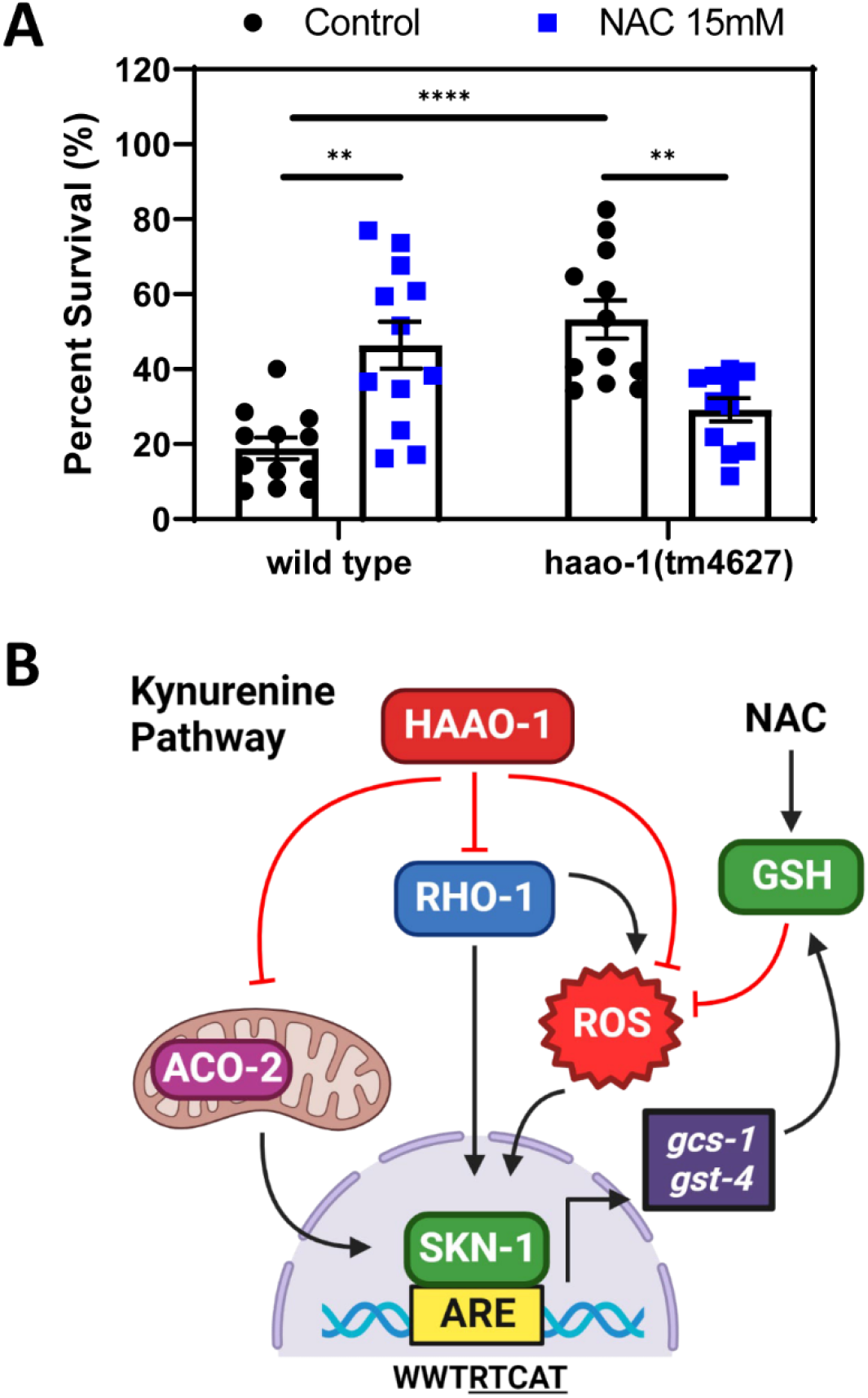
N-acetylcysteine (NAC) can diminish resistance to oxidative stress in haao-1(tm4627) background. (A) wild type worms grown from egg in NGM plates (n=456) or containing 15mM NAC (n=403), and *haao-1(tm4627)* worms grown from egg in NGM plates (n=444) or containing 15mM NAC (n=400). At L4 larval stage they were transferred to 200 μM juglone plates for 16 hours. Experimental animals were scored after 16 hours on juglone and counted as dead when not responding to prodding under a dissection microscope. (B) Schematic diagram of *haao-1* inhibition and *skn-1* upregulation by ROS-dependent and ROS-independent mechanisms involving *aco-2* and *rho-1*. P-values were determined by parametric unpaired t-test with Welch’s correction and using the Holm-Sidak multiple test correction method, with alpha = 0.05.

## Discussion

In this study, we report the effect of inhibition of the kynurenine pathway at the *haao-1* enzyme in the oxidative stress response in the nematode *C. elegans*. We determine that the enhanced response to several ROS-generating agents is dependent on the ability of haao-1 inhibition to induce ROS production and therefore activating the signaling pathways to respond to the oxidative damage in a hormetic fashion. We observed haao-1 knockdown promotes a better response to oxidative stress and activation of stress response pathways via several transcription factors as SKN-1, HIF-1, HSF-1 and DAF-16. Last, our study provides evidence that mitochondrial signaling (*aco-2*) and GTPase activity (*rho-1*) are mediators of SKN-1 induction in haao-1 knockout background.

Is likely that inhibition of the HAAO-1 enzyme increased the concentration of the upstream metabolite 3HAA (**Fig. 1A**) and this metabolite modifies the redox state of the cell by modulating the redox properties of the environment. Our previous findings demonstrate that haao-1 knockdown induces an increase in 3HAA to up to 10-fold while just moderately decrease NAD+ in worms (Dang et al., 2021).

There has been a lot of controversy related to the pro-oxidant or antioxidant properties of several metabolites in the kynurenine pathway and consequently, about their effects in the cell and potential roles in many redox-related processes(González Esquivel et al., 2017; Massudi et al., 2012; Sánchez Chapul et al., 2022; Sas et al., 2018; Wigner et al., 2017; Zhuravlev et al., 2016). However, recent evidence shows that the redox properties of many of these metabolites are dependent on the *in vivo* environment, particularly, the presence of other molecules around it. 3HAA, the main metabolite present in this study, has been shown to have pro-oxidant and antioxidant properties, but this effect also has been shown to be dependent on the presence of metal ions, concretely iron (Fe) (Chobot et al., 2015; Dang et al., 1998; Goldstein et al., 2000; Kubicova et al., 2013). And this effect is not just based on the need of iron as a co-factor to be able to carry on the enzymatic oxidative ring opening of the 3HAA by the HAAO-1 enzyme, but instead the modulation of the redox properties of the substrate by itself (Li, 2006; Wang et al., 2020; Yang et al., 2018).

One implication of our work is that modulation of the redox state of the cell through ROS generation by the accumulation of 3HAA inside of it will be an important consideration for future studies. These considerations actually point to very specific targeted disease contexts where ROS plays a crucial role. Cancer initiation and progression are two main particular scenarios where *haao-1* inhibition and 3HAA supplementation can be modulated to fundamentally turn a deleterious outcome into a more beneficial environment, from a medicinal perspective. During cancer initiation and progression, accumulation of ROS and subsequent DNA damage are two critical steps to maintain the mutation load for the malignant cells to acquire aggressive phenotypes. NAC, and FDA approved drug to prevent liver, lungs, and kidney damage, might be looked as a potential co-treatment for cancer patients with low HAAO protein expression.

Neurogenerative diseases are another branch where regulation of the redox state of the cell has an essential role. Specific somatic mutations in the catalytic domain of the HAAO enzyme might be able to cause accumulation of 3HAA in brain cells and change the delicate redox balance towards a more oxidative state and causing severe damage. From this perspective, modulation of several metabolites in the kynurenine pathway might have extensive repercussions in the development of several neurodegenerative disorders like Alzheimer’s disease (Wang et al., 2022), Parkinson’s disease (Heilman et al., 2020; Perez-Pardo et al., 2021), Huntington’s disease (Bondulich et al., 2021), and ALS (Fifita et al., 2021).

Together, this work provides evidence of an interesting mechanism of regulation for the antioxidant response by the redox properties of a specific endogenous metabolite in a highly conserved metabolic pathway. While previous work has been shown contradictory results about the redox properties and potential of the kynurenine pathway as an antioxidant or pro-oxidant modifier, this is the first evidence that shows in a model organism how the 3HAA metabolite can interact with its environment to induce ROS and prime the system for a better response to oxidative stress. Therefore, evidence shown here supports the model of 3HAA lifespan extension by priming the system with a hormetic effect based on ROS.

## Conclusions

By inhibiting the kynurenine pathway enzyme *haao-1*, we observed an overall increased response to oxidative stress. However, the increased activity of the transcription factor SKN-1/Nrf2 and its downstream target genes (*gcs-1, gst-4*, glutathione pathway) acts as a compensatory mechanism for the induced ROS production by *haao-1* inhibition and/or 3HAA redox properties. Overall, our results suggest that these interventions prime the redox state of the cell through a hormetic effect to be able to handle a secondary stressor in a more suitable manner. Our data supports the model that *haao-1* inhibition exerts a positive effect in the antioxidant response and promotes health span and increased lifespan.

## Notes

### Competing Interest Statement

The authors have declared no competing interest.

